# GenePT Revisited: Do Better Text Embeddings Make Better Gene Embeddings?

**DOI:** 10.64898/2026.04.16.718976

**Authors:** Jonathan G. Hedley, Philip H. S. Torr, Kaspar Märtens

## Abstract

GenePT introduced a simple recipe for gene representations: embed each gene’s natural-language description with a general-purpose text embedding model and reuse the resulting vectors across downstream tasks. Since GenePT’s release, embedding models have improved rapidly, with many strong open and commercial encoders benchmarked on suites such as the Massive Text Embedding Benchmark (MTEB). We present a controlled “leaderboard” study that keeps the GenePT pipeline fixed and varies only the embedding backbone. We benchmark contemporary encoders on four diverse gene embedding tasks: gene–gene interaction prediction, gene property classification, cell type classification, and prediction of transcriptomic responses to unseen genetic perturbations. Across these settings, newer backbones consistently outperform the original GenePT backbone (`text-embedding-ada-002`), achieving improvements of 1–17%, while enabling fully reproducible research by avoiding API dependencies.

## 1 Introduction

Learning useful gene representations is a recurring bottleneck in computational biology because many models need a compact, reusable way to encode gene identity and functional context, especially when generalizing beyond the genes, datasets, or conditions seen during training. While many approaches pretrain embeddings from large-scale omics data or protein sequence corpora, this can be expensive and often produces representations tied to a particular dataset or objective. Consequently, reusable gene embeddings are increasingly used as building blocks in larger systems; for example, single-cell “foundation” frameworks such as UCE (Rosen et al., 2024) and PULSAR (Pang et al., 2025) incorporate gene-level embeddings alongside expression to construct cellular representations.

GenePT (Chen & Zou, 2024) proposed a strikingly simple route to obtaining such embeddings. Each gene is represented by a natural-language description (e.g., curated NCBI summaries), which is then embedded with a general-purpose embedding model to yield a single vector per gene that can be reused across downstream tasks. This approach has been adopted broadly. scGenePT (Istrate et al., 2024) augments scGPT (Cui et al., 2024) by injecting GenePT-style language-derived gene embeddings, showing they provide a useful prior for learning cell-level models from expression. GenePT embeddings are also widely used as a prior for generalization tasks, most notably in *unseen perturbation prediction*, where they enable extrapolation beyond perturbations observed during training (Wang et al., 2024; Märtens et al., 2024; Ramakrishnan et al., 2025).

Since GenePT, the ecosystem of general-purpose embedding models has improved rapidly, with many strong open and commercial options available. This progress is reflected in broad, standardized benchmarks such as the Massive Text Embedding Benchmark (MTEB) (Enevoldsen et al., 2025). This raises a practical question for GenePT-style pipelines: *to what extent can we improve gene representations simply by swapping the underlying text embedding backbone, while keeping the rest of the pipeline fixed?* In this short “leaderboard-style” study, we answer this by isolating the effect of the embedding backbone and evaluate a panel of contemporary embedding models across four representative tasks that probe gene–gene relationships, gene-level labels, cell-level labels, and responses to unseen genetic perturbations. Across all experiments, we vary only the embedding backbone, holding the gene summaries, preprocessing, downstream modelling choices, and evaluation protocol constant. We find that newer backbones consistently improve performance over the GenePT baseline across these tasks, without domain-specific fine-tuning.

**Figure 1:**
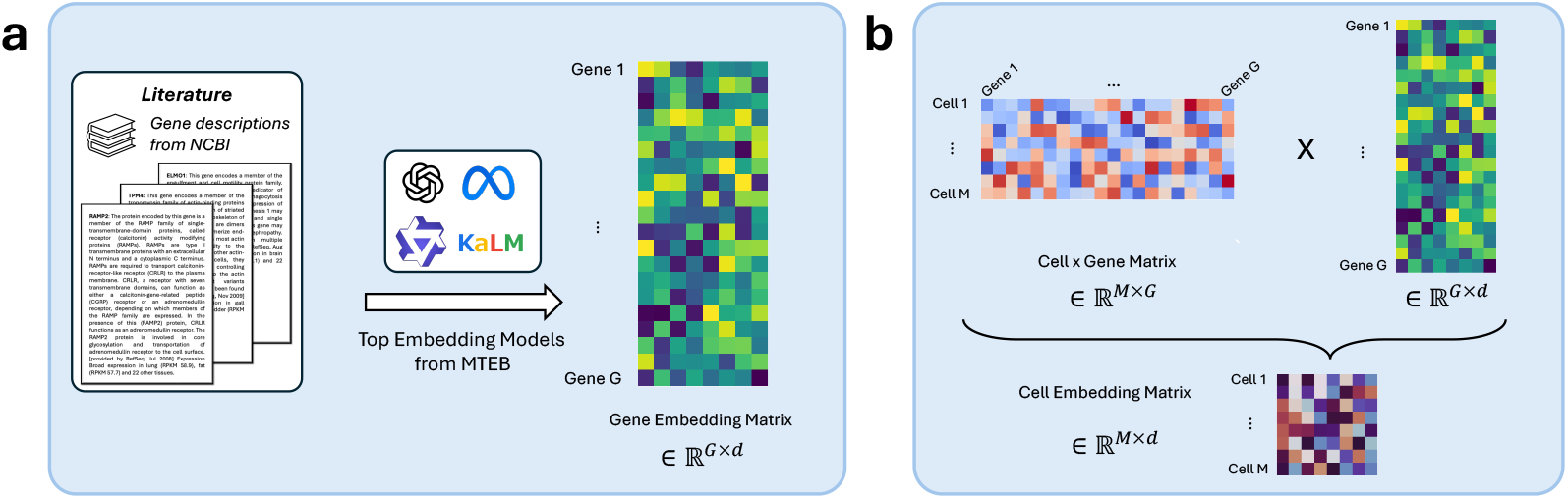
Overview. (a) We obtain a gene embedding matrix by feeding curated gene summaries through a chosen embedding model. (b) Cell embeddings are obtained by combining a cell-by-gene expression matrix with the gene embedding matrix.

## 2 Methods

### Gene text and gene embeddings

For each gene *g*, we embed a text description *t*_*g*_ (e.g. a curated summary) using an embedding model, *E*, yielding a gene vector **z**_*g*_ = *E*(*t*_*g*_) ∈ ℝ^*d*^, where *d* is the embedding dimension of *E*. To isolate the effect of the backbone, we keep the text source, formatting, and preprocessing fixed across all embedding models.

### Cell embeddings

For cell-level tasks, we derive cell embeddings from the gene embeddings. Let *X* ∈ ℝ^*M ×G*^ denote the (normalized) expression matrix for *M* cells and *G* genes, and let *Z* ∈ ℝ^*G×d*^ denote the gene embedding matrix. Following GenePT, we compute a cell embedding matrix *C* = *XZ* ∈ ℝ^*M ×d*^. Intuitively, each cell embedding is a weighted combination of gene vectors, with weights coming from the cell’s expression profile.

### Embedding Models

We compare the original GenePT backbone against several modern text embedding models selected from top-performing encoders on the Massive Text Embedding Benchmark (MTEB), which evaluates models across 9 task types (including retrieval, classification, clustering, and semantic similarity) spanning over 130 tasks and 250+ languages. We selected three top-ranking models: `KaLM-Embedding-Gemma3-12B-2511` (KaLM) (Zhao et al., 2025), `llama-embed-nemotron-8b` (Llama) (Babakhin et al., 2025), and `Qwen3-Embedding-8B` (Qwen3) (Zhang et al., 2025). All three are open-weight models trained on general web corpora via multi-stage contrastive learning, without targeted biological optimization. This makes performance gains attributable to improved general-purpose text understanding rather than domain adaptation. In contrast to GenePT’s reliance on the proprietary `text-embedding-ada-002` API, these models enable fully reproducible research with local deployment^1^.

We also include protein language model baselines ProtT5 (Elnaggar et al., 2022) and ESM2 (Lin et al., 2023). Across experiments, only the embedding backbone varies; everything else is fixed.

## 3 Experiments

### 3.1 Task 1: Gene-Gene Interaction Prediction

We first test whether gene embeddings capture functional relationships by predicting gene–gene interactions (GGIs). Each example is a pair of genes (*g*_*i*_, *g*_*j*_) labeled as interacting or not, using a benchmark for GGIs derived from shared gene ontology annotations (Du et al., 2019). We therefore represent each fixed gene pair by the summed embedding 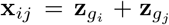, on which we train a binary classifier and evaluate via ROC-AUC (curves in Appendix A; values in Table 3).

Across both logistic regression (LR) and random forest (RF) classifiers, modern text embedding backbones consistently outperform the GenePT baseline, while protein-sequence embeddings lag behind in this text-derived benchmark. Among text models, Llama achieves the strongest performance, improving over GenePT by +2.9% (LR) and +0.9% (RF) in ROC-AUC. Qwen3 is close behind, with performance differences narrowing under the stronger random forest classifier. As shown in the ROC curves (Appendix A), the Gene2Vec baseline remains substantially weaker, and random embeddings perform at chance, confirming that the task is sensitive to representation quality rather than the classifier.

**Table 1:**
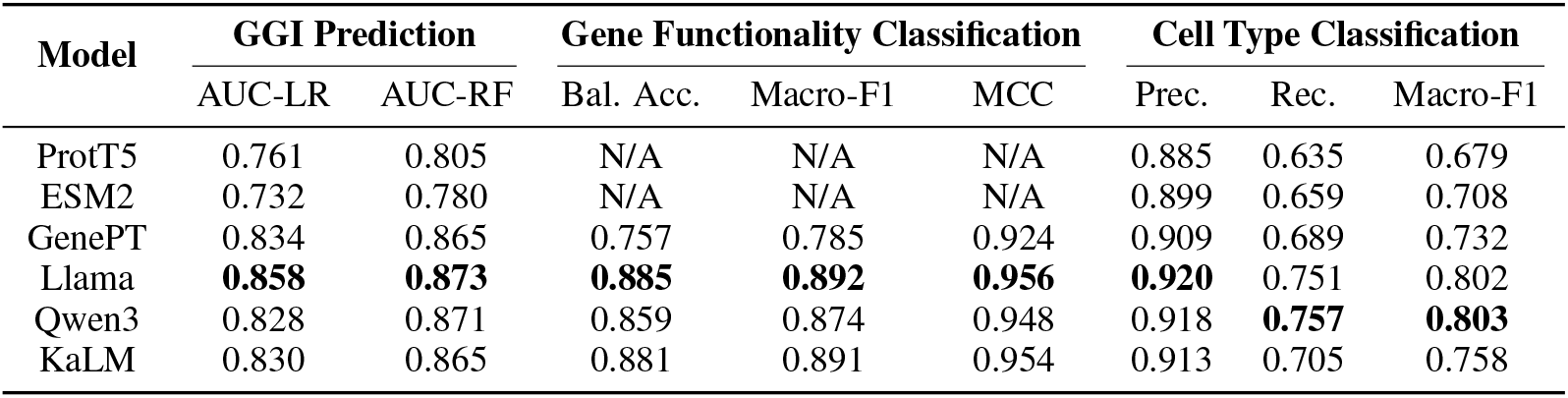
Summary performance across three tasks. We report gene-gene interaction prediction (ROC-AUC under logistic regression (LR) and random forest (RF)), gene property classification (balanced accuracy, macro-F1, and Matthews Correlation Coefficient), and cell type classification (precision/recall/macro-F1). Higher is better for all metrics shown. Modern text embedding backbones consistently outperform GenePT embeddings and protein-sequence baselines.

### 3.2 Task 2: Gene Functionality Classification

Here we evaluate gene functionality classification, a multi-class task assigning each gene to a categorical label (e.g., functional class). For each embedding backbone, we train a simple classifier on the corresponding gene embeddings and report balanced accuracy, macro-F1, and Matthews correlation coefficient (MCC). Balanced accuracy averages per-class recall and is less sensitive to class imbalance than standard accuracy; macro-F1 averages the per-class F1 scores, weighting rare classes equally; and MCC provides a single summary of prediction quality that accounts for all entries of the confusion matrix.

Modern text embedding backbones substantially improve class separation relative to earlier text encoders. This is reflected in the summary metrics and in the normalized confusion matrices (Appendix B), where stronger backbones yield a cleaner diagonal and reduce systematic confusions between related classes. Llama performs best overall (Table 3), improving over GenePT by +16.9% in balanced accuracy, +13.6% in macro-F1, and +3.5% in MCC. Results for ProtT5 and ESM2 are omitted here because their embeddings are only available for a subset of protein-coding genes, which does not match the gene set used in this task.

### 3.3 Task 3: Cell Type Classification

Next, we evaluate whether gene embeddings support cell identity discrimination in a single-cell setting, using cells taken from the Aorta dataset (Li et al., 2020) (11 cell types). We construct cell-level representations using the expression-weighted aggregation described in Sec. 2. We then assess whether cells of the same annotated type cluster together by training a kNN classifier in the resulting cell embedding space, using an 80/20 train-test split. We report accuracy and macro-averaged precision/recall/F1, again emphasizing macro-F1 as a robust summary under class imbalance.

Echoing the previous tasks, modern text embedding backbones yield substantially better cell-type separability than the GenePT baseline and outperform protein-language-model baselines in this setting (Table 3). Relative to GenePT, Qwen3 improves recall by +9.9% and macro-F1 by +9.7%, while Llama achieves the strongest improvement in precision of +1.2%. KaLM improves over GenePT but remains behind Llama and Qwen3 overall. Together, these gains indicate more balanced performance across cell types under class imbalance.

### 3.4 Task 4: Genetic Perturbation Prediction

Finally, we evaluate genetic perturbation response prediction, where the goal is to generalize to unseen perturbations using a regressor conditioned on gene embeddings. Given a genetic perturbation *p* and its corresponding text embedding **z**_*p*_ ∈ ℝ^*d*^, we train an MLP *f*_*θ*_ : ℝ^*d*^ → ℝ^*G*^ to predict pergene mean expression shifts Δ*µ* = *µ*_*p*_− *µ*_ctrl_, where *µ*_*p*_ ∈ ℝ^*G*^ denotes the vector of mean gene expression across cells under perturbation *p* and *µ*_ctrl_ is the control mean. This setup is an established approach for generalizing perturbation prediction models and has been shown to outperform single-cell foundation model baselines on this task (Märtens et al., 2024; Ramakrishnan et al., 2025).

We evaluate performance using (i) Pearson correlation between predicted and observed mean shifts over the top *K* differentially expressed genes and (ii) relative mean absolute error (RMAE) computed over all genes, reporting mean ± s.d. over 9-fold cross-validation in Table 2. Across datasets and cell lines, text embedding backbones consistently outperform protein language model baselines. Llama is strongest overall: on K562 it improves over GenePT by roughly +3–4% in correlation and reduces RMAE by ∼1.6%. On RPE1, Llama gives small but consistent gains over GenePT on larger DE sets (Top 50/100) and lowers RMAE by ∼1.5%, while KaLM is best on the smallest DE set (Top 20). On HEPG2, GenePT remains best on Top 20, but Llama achieves the best Top 50/100 correlations and reduces RMAE by ∼1.1%.

**Table 2:** Perturbation response prediction on unseen perturbations across multiple cell lines. We report mean prediction performance (± s.d. across folds) on unseen perturbations from the K562 and RPE cell lines from Replogle et al. (2022) and the HEPG2 cell line from Nadig et al. (2025). We report Pearson correlations over the top *K* ∈ *{*20, 50, 100*}* DE genes and RMAE over all genes.

| Cell line | Embedding Model | Pearson correlation (↑) | | | RMAE (↓) |
| --- | --- | --- | --- | --- | --- |
|  |  | Top 20 | Top 50 | Top 100 |  |
| K562 | ProtT5 | 0.579 ± 0.020 | 0.625 ± 0.014 | 0.635 ± 0.023 | 0.917 ± 0.018 |
| | ESM2 | 0.570 ± 0.041 | 0.610 ± 0.033 | 0.619 ± 0.028 | 0.925 ± 0.020 |
| | GenePT | 0.644 ± 0.048 | 0.685 ± 0.041 | 0.691 ± 0.038 | 0.886 ± 0.029 |
|  | Llama | **0.670 ± 0.035** | **0.710 ± 0.028** | **0.716 ± 0.026** | **0.872 ± 0.029** |
| | Qwen3 | 0.660 ± 0.041 | 0.705 ± 0.035 | 0.711 ± 0.033 | 0.875 ± 0.028 |
| | KaLM | 0.649 ± 0.044 | 0.693 ± 0.034 | 0.699 ± 0.033 | 0.879 ± 0.025 |
| RPE1 | ProtT5 | 0.665 ± 0.030 | 0.713 ± 0.027 | 0.733 ± 0.029 | 0.893 ± 0.047 |
| | ESM2 | 0.662 ± 0.026 | 0.709 ± 0.026 | 0.728 ± 0.028 | 0.897 ± 0.039 |
| | GenePT | 0.682 ± 0.031 | 0.736 ± 0.033 | 0.758 ± 0.031 | 0.873 ± 0.040 |
| | Llama | 0.688 ± 0.030 | **0.740 ± 0.030** | **0.762 ± 0.028** | **0.860 ± 0.034** |
| | Qwen3 | 0.687 ± 0.028 | 0.738 ± 0.030 | 0.761 ± 0.027 | 0.865 ± 0.044 |
| | KaLM | **0.690 ± 0.035** | 0.738 ± 0.035 | 0.760 ± 0.032 | 0.864 ± 0.041 |
| HEPG2 | ProtT5 | 0.438 ± 0.019 | 0.454 ± 0.023 | 0.449 ± 0.024 | 0.981 ± 0.019 |
| | ESM2 | 0.421 ± 0.024 | 0.437 ± 0.023 | 0.430 ± 0.021 | 0.990 ± 0.024 |
| | GenePT | **0.485 ± 0.039** | 0.486 ± 0.032 | 0.481 ± 0.028 | 0.969 ± 0.021 |
| | Llama | 0.477 ± 0.031 | **0.493 ± 0.027** | **0.487 ± 0.027** | **0.958 ± 0.022** |
| | Qwen3 | 0.484 ± 0.030 | 0.492 ± 0.027 | 0.485 ± 0.027 | 0.961 ± 0.023 |
| | KaLM | 0.476 ± 0.035 | 0.484 ± 0.030 | 0.477 ± 0.029 | 0.969 ± 0.020 |

## 4 Conclusion

Across gene-gene interaction, gene classification, cell type classification, and perturbation prediction tasks, replacing the GenePT embedding backbone for stronger general-purpose text embedding models yields consistent improvements of 1–17%, depending on task and metric. These gains require no biological pretraining and no changes to the downstream pipeline. Although newer models often produce higher-dimensional embeddings, the improvements are not simply a consequence of embedding size: in matched-dimension comparisons using PCA for Tasks 1–3, the overall advantage of the newer language models is broadly preserved (Appendix C), suggesting that the gains are driven primarily by better representation quality rather than dimensionality alone. Moreover, unlike GenePT’s reliance on proprietary APIs, these open-weight models enable fully reproducible research with local deployment.

We have released pre-computed gene embeddings for all evaluated models, eliminating computational barriers to adoption. Moreover, as even stronger text embedding models emerge on the MTEB leaderboard, GenePT-style pipelines stand to benefit automatically. Our results suggest that improvements in general-purpose text understanding translate directly to better gene representations.

While we find that top-performing MTEB encoders apply well to gene descriptions, these models are not trained specifically for biological text. Future gains will likely come from both stronger encoders and improved gene descriptions: for instance, standardized formats highlighting function and pathway context, or hybrid approaches enriching sparse text with structured annotations. Overall, for practitioners using GenePT-style pipelines today, switching to top-performing open-weight MTEB models provides immediate benefits at minimal implementation cost.

## Acknowledgments

JGH acknowledges support from a Novo Nordisk Postdoctoral Fellowship in partnership with the University of Oxford, and the support of the David Cockayne Junior Research Fellowship from Linacre College, Oxford.

## Appendix

### A Gene-Gene Interaction Prediction: ROC Curves

**Figure 2:**
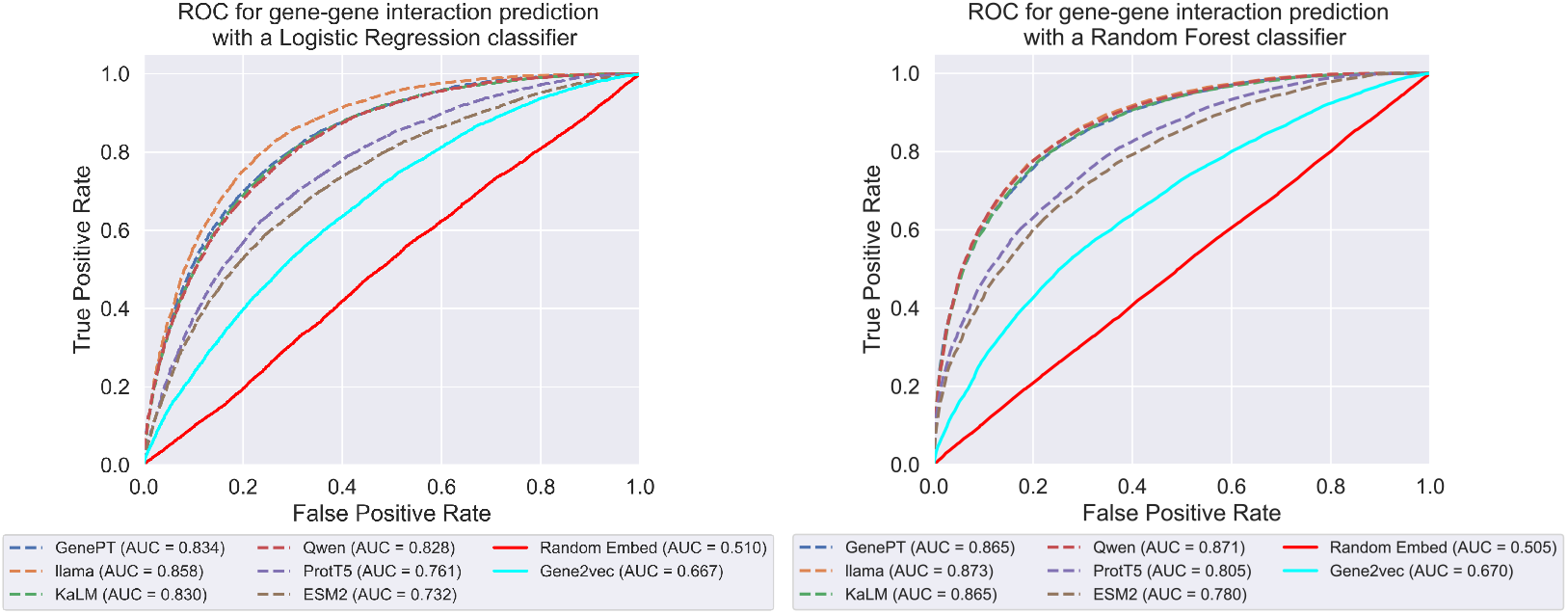
ROC curves for gene-gene interaction prediction under simple classification models, illustrating consistent gains from newer text embedding backbones over GenePT and biological baselines.

### B Gene Functionality Classification: Confusion Matrices

**Figure 3:**
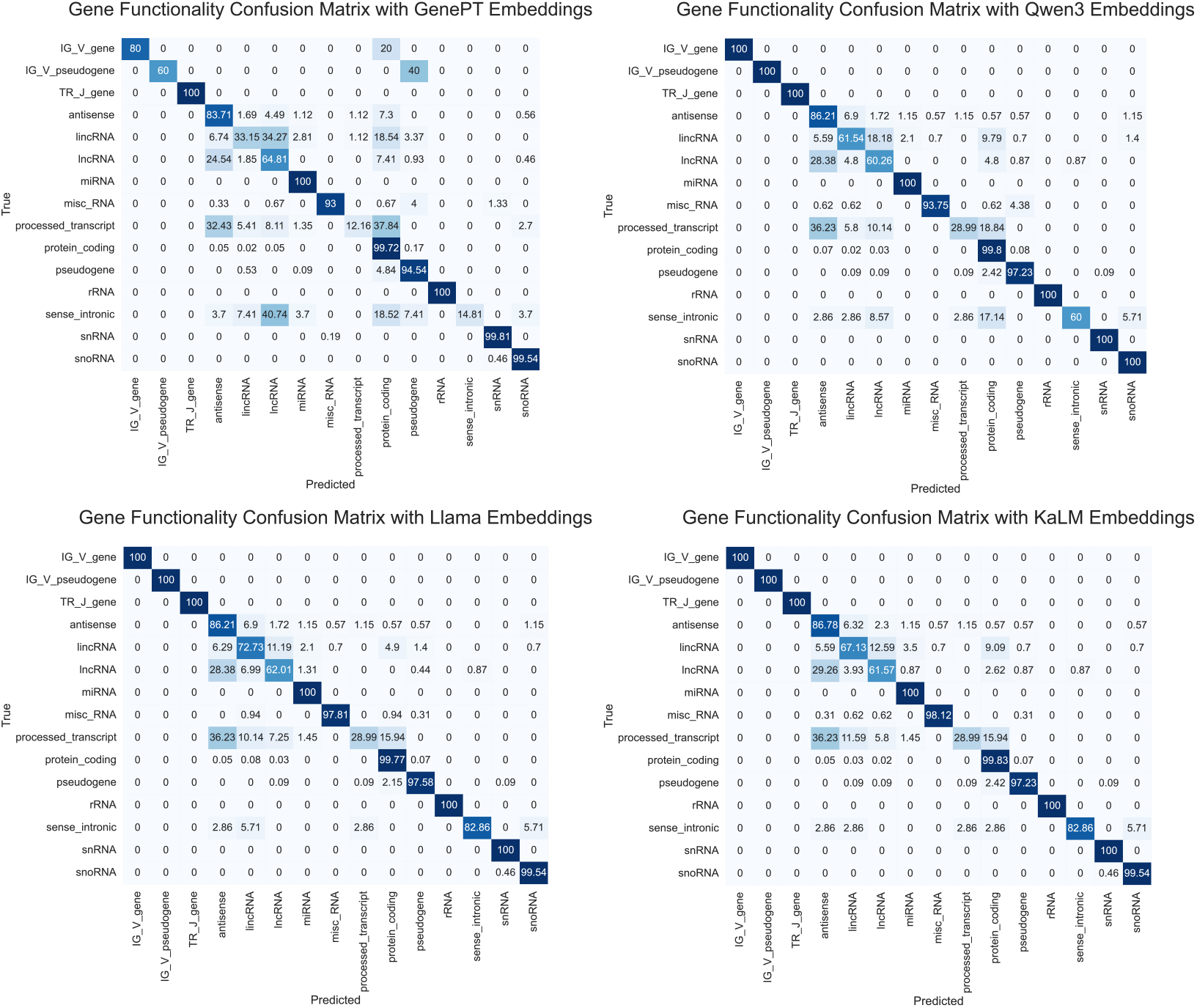
Normalized confusion matrices for gene property classification comparing GenePT embeddings with modern embedding models, showing overall improved class separation and fewer systematic confusions.

### C Results with PCA-Matched Dimensions

For each train-test split, PCA was fitted on the training embeddings only and then applied to the corresponding test embeddings, to avoid information leakage.

**Table 3:** Summary performance across three tasks. We report gene-gene interaction prediction (ROC-AUC under logistic regression (LR) and random forest (RF)), gene functionality classification (balanced accuracy, macro-F1, and Matthews Correlation Coefficient), and cell type classification (precision/recall/macro-F1). Higher is better for all metrics shown. For each model, we report results at its native embedding dimensionality, as well as after PCA reduction to matched lower-dimensional representations. Modern text embedding backbones consistently outperform GenePT embeddings and protein-sequence baselines, and these overall trends remain broadly preserved under PCA-based dimensionality control.

| Model | Dim. | GGI Prediction |  | Gene Functionality Classification |  |  | Cell Type Classification |  |  |
| --- | --- | --- | --- | --- | --- | --- | --- | --- | --- |
|  |  | AUC-LR | AUC-RF | Bal. Acc. | Macro-F1 | MCC | Prec. | Rec. | Macro-F1 |
| ProtT5 | 1024 | 0.758 | 0.805 | N/A | N/A | N/A | 0.885 | 0.635 | 0.679 |
| ESM2 | 5120 | 0.732 | 0.780 | N/A | N/A | N/A | 0.899 | 0.659 | 0.708 |
| GenePT | 1536 | 0.834 | 0.865 | 0.757 | 0.785 | 0.924 | 0.909 | 0.689 | 0.732 |
| Llama | 4096 | **0.858** | **0.873** | **0.885** | **0.892** | **0.956** | **0.920** | 0.751 | 0.802 |
| Qwen3 | 4096 | 0.828 | 0.871 | 0.859 | 0.874 | 0.948 | 0.918 | **0.757** | **0.803** |
| KaLM | 3840 | 0.830 | 0.865 | 0.881 | 0.891 | 0.954 | 0.913 | 0.705 | 0.758 |
| ProtT5 | 1024 | 0.758 | 0.805 | N/A | N/A | N/A | **0.885** | 0.635 | 0.679 |
| ESM2 | 1024 | 0.755 | 0.747 | N/A | N/A | N/A | 0.849 | 0.639 | 0.661 |
| GenePT | 1024 | 0.836 | 0.788 | 0.761 | 0.795 | 0.927 | 0.861 | 0.689 | 0.720 |
| Llama | 1024 | **0.859** | **0.854** | **0.870** | **0.876** | **0.952** | 0.866 | 0.688 | 0.713 |
| Qwen3 | 1024 | 0.829 | 0.838 | 0.839 | 0.856 | 0.943 | 0.862 | **0.709** | **0.729** |
| KaLM | 1024 | 0.829 | 0.800 | 0.861 | 0.874 | 0.950 | 0.860 | 0.694 | 0.708 |
| ProtT5 | 128 | 0.773 | 0.800 | N/A | N/A | N/A | 0.723 | 0.606 | 0.616 |
| ESM2 | 128 | 0.762 | 0.780 | N/A | N/A | N/A | 0.775 | 0.638 | 0.660 |
| GenePT | 128 | 0.813 | 0.860 | 0.729 | 0.747 | 0.919 | 0.849 | 0.676 | 0.707 |
| Llama | 128 | **0.836** | **0.882** | 0.830 | 0.843 | 0.941 | **0.885** | **0.698** | **0.738** |
| Qwen3 | 128 | 0.823 | 0.877 | 0.808 | 0.826 | 0.937 | 0.872 | 0.688 | 0.722 |
| KaLM | 128 | 0.822 | 0.859 | **0.844** | **0.858** | **0.944** | 0.784 | 0.675 | 0.692 |

## Footnotes

1 Pre-computed gene embeddings for all models, with code to reproduce our results are available at https://github.com/jghedley/gene_embedding_leaderboard

